# Critical dynamics in spontaneous EEG predict perturbational complexity in disorders of consciousness with measurable evoked responses

**DOI:** 10.64898/2026.06.04.730221

**Authors:** Derek Newman, Charlotte Maschke, Jordan O’Byrne, Michele Angelo Colombo, Angela Comanducci, Silvia Casarotto, Giuseppe Citerio, Mario Rosanova, Marcello Massimini, Karim Jerbi, Stefanie Blain-Moraes

**Affiliations:** Montreal General Hospital, McGill University Health Centre, Montreal, Canada; Integrated Program in Neuroscience, McGill University, Montreal, Canada; Aix-Marseille Université, Inserm, Institut de Neurosciences des Systèmes, UMR 1106, Marseille, France; Psychology Department, University of Montreal, Montreal, QC, Canada; MILA (Québec Artificial Intelligence Institute), Montréal, QC, Canada; Department of Biomedical and Clinical Sciences, University of Milan, Milan 20157, Italy; IRCCS, Fondazione Don Carlo Gnocchi ETS, Milan 20148, Italy; Scuola di Medicina e Chirurgia, University of Milan Bicocca, Milan, 20126, Italy; Centre UNIQUE (Union Neurosciences & Intelligence Artificielle), Montréal, QC, Canada; Department of Occupational Sciences and Occupational Therapy, University of British Columbia, Vancouver, Canada; School of Biomedical Engineering, University of British Columbia, Vancouver, British Columbia, Canada

**Keywords:** neural criticality, Perturbational Complexity Index, disorders of consciousness, severe brain injury, electroencephalography, spontaneous brain dynamics

## Abstract

Identifying which severely brain-injured patients retain the capacity for consciousness remains a major challenge in neurocritical care. The perturbational complexity index (PCI) provides a reliable assessment of consciousness capacity, but its reliance on transcranial magnetic stimulation and EEG (TMS-EEG) limits bedside scalability. PCI and brain criticality capture complementary dimensions of brain dynamics: PCI quantifies the complexity of the brain’s evoked response to perturbation, whereas criticality characterizes the intrinsic organization of spontaneous activity. Here, we tested whether resting-state EEG signatures of criticality predict PCI_max_ in disorders of consciousness, extending prior findings from anesthesia to severe brain injury. In 26 patients with vascular, traumatic, or anoxic brain injury, multivariate criticality related features did not generalize PCI_max_ prediction across the full heterogeneous cohort. However, criticality features predicted PCI_max_ when analyses were restricted to non-anoxic patients and when restricting analyses to patients with non-zero PCI_max_ values. These findings suggest that spontaneous criticality measures index the brain’s intrinsic dynamical regime that supports complex perturbational responses, while their correspondence with PCI_max_ depends on whether the injured brain retains sufficient capacity to sustain large-scale evoked responses. Together, our results extend the relationship between resting-state criticality and evoked perturbational complexity to disorders of consciousness and support the development of stratified EEG measures in severe brain injury.

## Introduction

Determining whether a severely brain-injured patient retains the capacity for consciousness remains one of the most consequential challenges in clinical neuroscience and neurocritical care. In disorders of consciousness (DoC), overt behavior is often minimal or absent, yet behavioral unresponsiveness does not necessarily imply the absence of conscious capacity. This uncertainty has major ethical and clinical implications, shaping diagnosis, prognosis, pain management, treatment decisions, and communication with families^1^. The central problem is therefore not only to detect behavior, but to infer capacity of the injured brain to sustain consciousness.

The Perturbational Complexity Index (PCI) has emerged as a promising tool for addressing this problem. PCI quantifies the spatiotemporal complexity of cortical responses evoked by transcranial magnetic stimulation (TMS), capturing the presence of large-scale integration and differentiation in brain activity^2^. By probing how the brain responds to direct cortical perturbation, PCI provides an assessment of cortico-thalamo-cortico interactions independently of sensory, processing, executive function and motor behaviour. In benchmark conditions, including wakefulness, sleep, dreaming and anesthesia, PCI has shown high accuracy in distinguishing conscious from unconscious conditions^3^, which allows inferring a covert capacity for consciousness in patients who remain unresponsive after brain injury^4^. Yet despite its conceptual power, PCI remains difficult to deploy widely in routine care because it requires specialized equipment and advanced expertise, limiting its bedside scalability^5,6^. A passive resting-state EEG proxy for PCI would therefore fill an important translational gap between mechanistic consciousness science and clinical practice.

One promising framework for such a proxy comes from the hypothesis that the brain operates near criticality. Critical systems are poised near a transition regime between excessive order and disorder^7^ that maximizes dynamic range^8^, promotes information transmission and capacity^9^, and supports flexible yet stable responses to perturbation^10^. In neuroscience, criticality has been linked to metastability, scale-free activity, avalanche dynamics, long-range temporal correlations observed across brain states^11,10,7,12^ and levels of consciousness^13,14,15^. Although debate remains regarding how uniformly or literally criticality should be interpreted in brain systems^16^, the framework is increasingly viewed as clinically relevant because it offers a mechanistic account for describing how intrinsic neural dynamics may reflect responses to stimulation and conscious state transitions^17–19,15^.

This framework is especially compelling in the context of consciousness research. Complexity-based accounts have long emphasized that conscious states are distinguished not by maximal activity, but by the capacity to sustain richly differentiated yet integrated dynamics^20–22^. In that sense, criticality and perturbational complexity are deeply related: criticality describes an intrinsic dynamical regime favorable to flexible large-scale interactions, whereas perturbational complexity measures evoked response elicited when that system is externally perturbed. Recent work from our group demonstrated that in healthy participants undergoing anesthesia, resting-state EEG criticality features predicted perturbational complexity, suggesting that the brain’s intrinsic dynamical state constrains its capacity to generate complex responses to stimulation^14^.

However, whether this relationship extends to severely brain-injured patients remains unknown and may not be expected to hold uniformly across all forms of severe brain injury. PCI is only informative when TMS evokes a measurable spatiotemporally distributed response. When cortical responses are absent, highly localized or suppressed, PCI collapses toward a floor value, reflecting a failure of propagation. The mapping between spontaneous criticality and PCI may be conditional on a preserved perturbational response regime. Unlike pharmacologically altered states in non-injured brains, DoC arise in the setting of heterogeneous structural damage, disrupted network architecture, and large inter-individual variability. Specifically, anoxic injuries may produce profoundly abnormal EEG activity and reduced cortical responsiveness, potentially weakening or distorting relationships that hold in healthier brains^23,24^.

Here, we test whether resting-state EEG criticality can predict perturbational complexity in a clinical population of patients with disorders of consciousness. Specifically, we asked whether subject-level criticality features extracted from spontaneous EEG could predict PCI_max_, the maximum PCI value observed across stimulation sites. Our hypothesis is illustrated in Figure 1 (left). Since PCI depends on the capacity of the cortex to generate measurable EEG response to TMS, we further examined whether the criticality-PCI relationship differed in patients with reduced or absent cortical responsiveness. We first assessed this with particular attention to etiology, comparing anoxic versus non-anoxic injuries, and then examined patients with absent perturbational complexity, defined by PCI_max_= 0. We also characterized the multivariate organization of the criticality feature space and examined univariate feature–PCI_max_ relationships across etiologies. Finally, to assess clinical relevance, we performed post-hoc analyses linking EEG criticality features and their low-dimensional latent structure to behavioural responsiveness (CRS-R)^25^ and functional outcome (GOS-E)^26^. By testing these hypotheses in a heterogeneous DoC cohort, we aimed to determine whether passive EEG criticality can serve as a clinically tractable marker of the brain’s capacity for consciousness.

**Figure 1.**
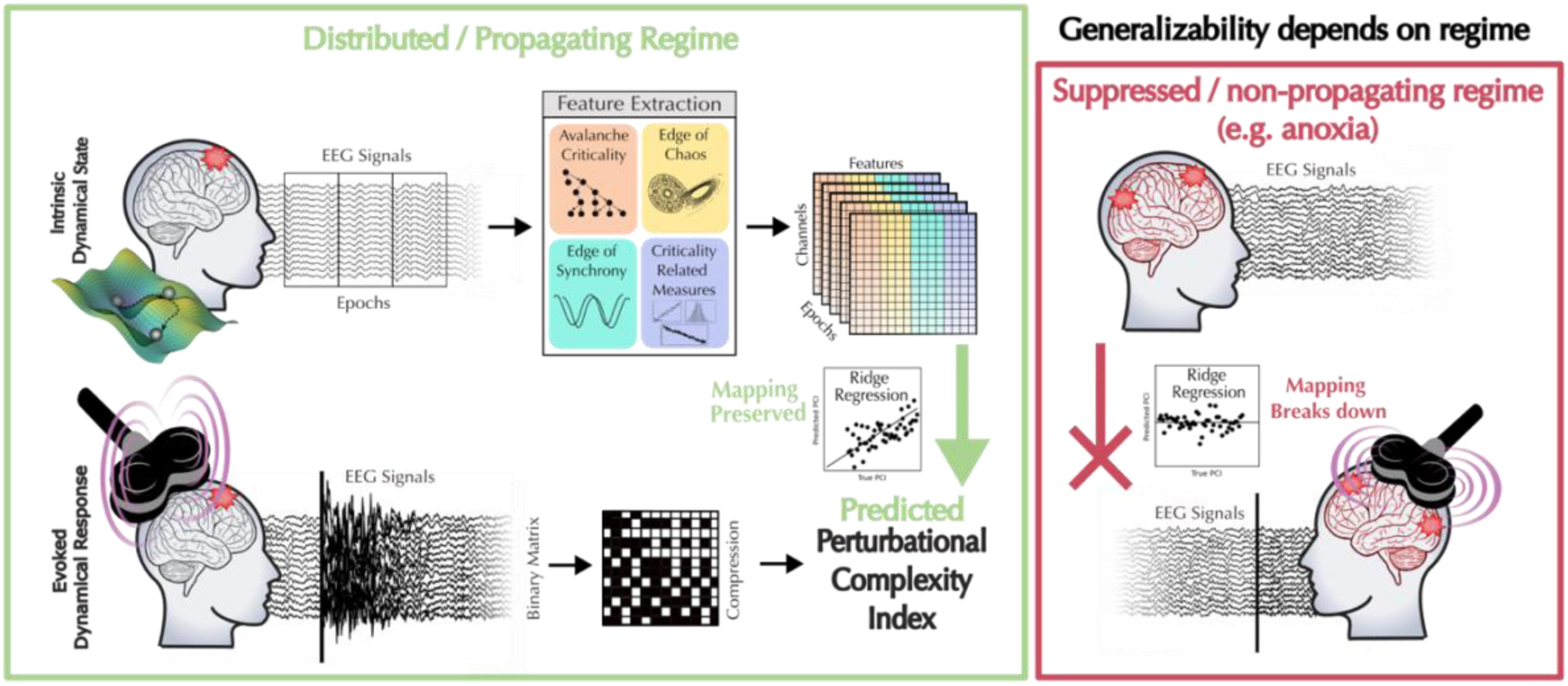
Regime-dependent prediction of perturbational complexity from spontaneous EEG criticality in patients in disorders of consciousness. Resting-state EEG of patients in disorders of consciousness was preprocessed, segmented into epochs, and used to extract multivariate features spanning avalanche criticality, which captures whether cascades of neural activity propagate with scale-free event statistics; edge-of-chaos dynamics, which quantify whether neural activity is stable, unstable, or sensitive to initial conditions; edge-of-synchrony, which measures the balance between excessive synchronous and asynchronous neural activity; and additional measures related to criticality, including temporal scaling, fractality and complexity, which quantify long-range temporal correlations, irregularity, and structured dynamical patterns. These features are organized across channels and epochs to form a multivariate feature matrix describing each patient’s spontaneous EEG dynamics. A ridge regression model is trained to predict the maximal Perturbational Complexity Index values (PCI_*max*_), derived independently from TMS-EEG evoked responses via spatiotemporal binarization and compression. This overview highlights that this prediction is conditional on the evoked cortical response regime. In the distributed/propagating regime (left; green box), TMS evokes a measurable, spatially distributed cortical response, resulting in non-zero PCI values. In this regime, spontaneous EEG criticality features can be meaningfully mapped onto the evoked perturbational complexity. In the suppressed/non-propagating regime (right; red box), TMS fails to evoke a distributed response and PCI collapses toward a floor value of zero. In this case, the absence of evoked propagation creates a mapping problem; spontaneous EEG features cannot reliably predict PCI because the target variable reflects floor level responses rather than graded perturbational complexity. Thus, including patients with absent TMS propagation (e.g. patients with severe anoxic brain injury) reduces generalizability by mapping spontaneous criticality features with non-informative PCI values.

## Results

We analyzed data from previously published studies of brain-injured patients with various progressions of disorders of consciousness^4,23^ consisting of three primary etiologies: vascular injuries (*n* = 13), anoxic injuries (*n* = 9) and traumatic injuries (*n* = 4). Resting-state electroencephalography (EEG) was recorded in the clinical environment contextually to the PCI protocol. PCI_*max*_ was assessed using a TMS-EEG protocol^2^. Prior to EEG acquisition, all patients had been off sedative medication for at least seven days^4,23^.

### Baseline criticality does not predict PCI_*max*_ across all DoC etiologies when floor-level non-responses are pooled with measurable responses

We first examined whether spontaneous EEG criticality features predict perturbational complexity across the full patient cohort (*n* = 26), irrespective of etiology. Using ridge regression trained on 16 criticality features (see Methods), we modeled PCI_*max*_ in all 26 patients and assessed performance using both in-sample prediction and leave-one-out cross-validation (LOO-CV). In-sample performance reflects the strength of association and variance explained within the analyzed cohort, whereas LOO-CV evaluates whether this relationship generalizes to an unseen patient and reflects the predictive value.

Across the full cohort, when all etiologies are analyzed together, the model shows a moderate in-sample fit (R² = 0.29), with predicted PCI_*max*_ values correlated with the true values (r = 0.60, p = 0.001). In-sample prediction error was modest, with a mean squared error (MSE) of 0.02 and a mean absolute error (MAE) of 0.11, corresponding to an average absolute prediction deviation of approximately 0.11 PCI_*max*_ units. However, this relationship does not generalize across unseen patients. Under LOO-CV, predictive performance collapses, yielding a negative R² (–0.13), increased prediction error (MSE = 0.032; MAE = 0.139), and no significant association between predicted and observed PCI_*max*_ (r = -0.046, p = 0.824) (Fig. 2).

**Figure 2.**
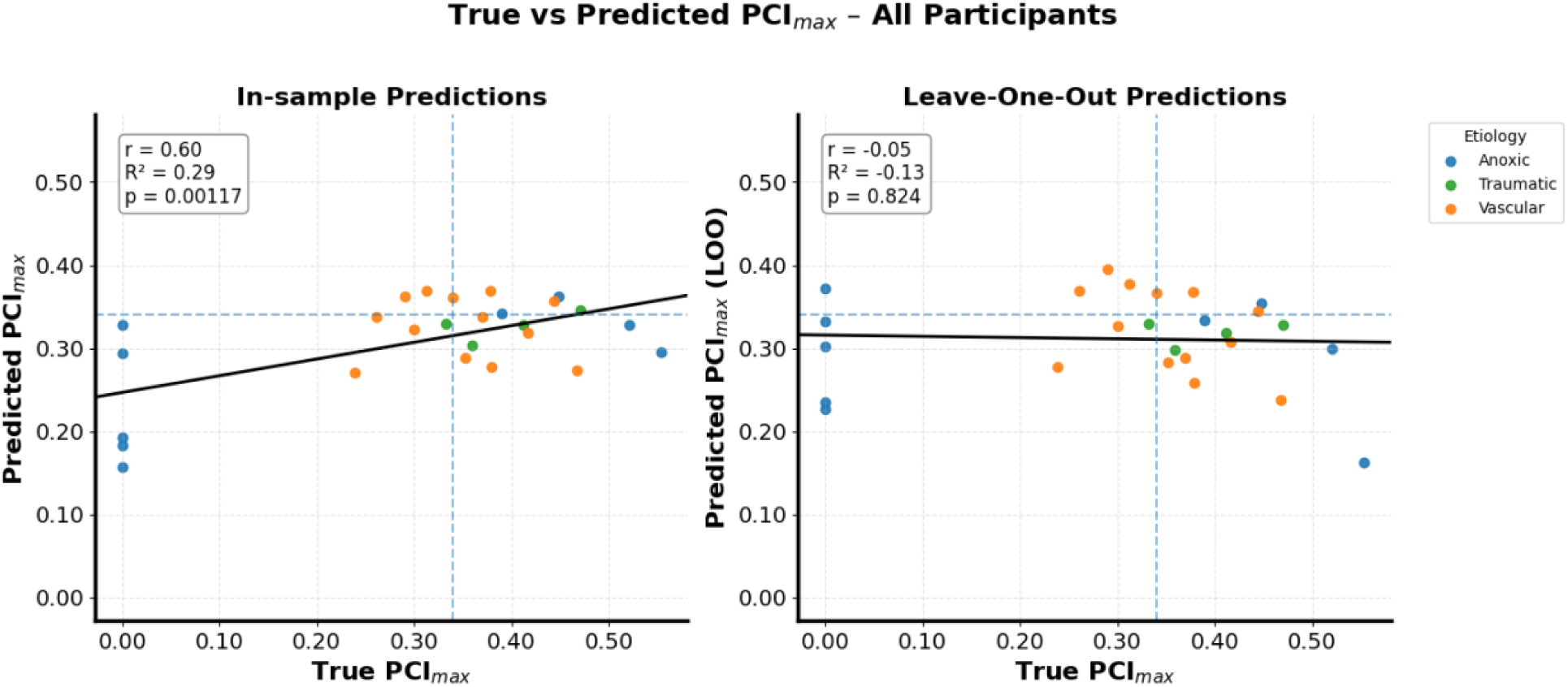
Baseline EEG criticality does not generalize to perturbational complexity when etiologies are pooled. True versus predicted PCI_*max*_ values obtained from a ridge regression model trained on 16 resting-state EEG criticality features across all patients (n = 26). **Left**, in-sample predictions show a moderate apparent association between predicted and observed PCI_*max*_ values. **Right**, leave-one-out cross-validation (LOO-CV) predictions reveal a complete loss of generalization, with near-zero correlation. Points are colored by injury etiology (anoxic-blue; traumatic-green; vascular-orange). Dashed lines indicate the empirical PCI threshold (PCI = 0.34). Together, these results demonstrate that when injury etiologies are combined, patient heterogeneity obscures any reliable mapping between intrinsic critical dynamics and evoked perturbational complexity.

These results indicate that when etiologies are pooled, baseline spontaneous criticality features do not reliably predict PCI_*max*_ across patients. While the model captures a modest relationship within the training data, this mapping is unstable across patients as criticality features fail to generalize under LOO-CV, indicating poor prediction of PCI_*max*_ for individual unseen data. The inclusion of floor-level PCI_*max*_ responses likely obscure the relationship between intrinsic spontaneous dynamics and the capacity to generate complex evoked responses to perturbation.

### Baseline criticality predicts PCI_*max*_ in non-anoxic DoC patients

Given that anoxic brain-injury is often characterized by abnormal EEG activity, reduced cortical responsiveness, and diffuse disruption of large-scale cortical organization^23^, we next restricted the analysis to non-anoxic patients (vascular and traumatic etiologies; *n* = 17), a subgroup expected to retain relatively more preserved large-scale cortical organization than anoxic injury.

The in-sample model demonstrated a strong fit (R² = 0.66), explaining 66% of the variance in PCI_*max*_ within the training data, with predicted values closely tracking empirical measurements (r = 0.812, p < 0.001). Importantly, this relationship generalized across unseen patients. Under LOO-CV, the model retained substantial predictive power (R² = 0.365), with a low prediction error (MSE = 0.0028; MAE = 0.044) and a significant correlation between predicted and observed PCI_*max*_ (r = 0.634, p = 0.0063) (Fig. 3).

**Figure 3.**
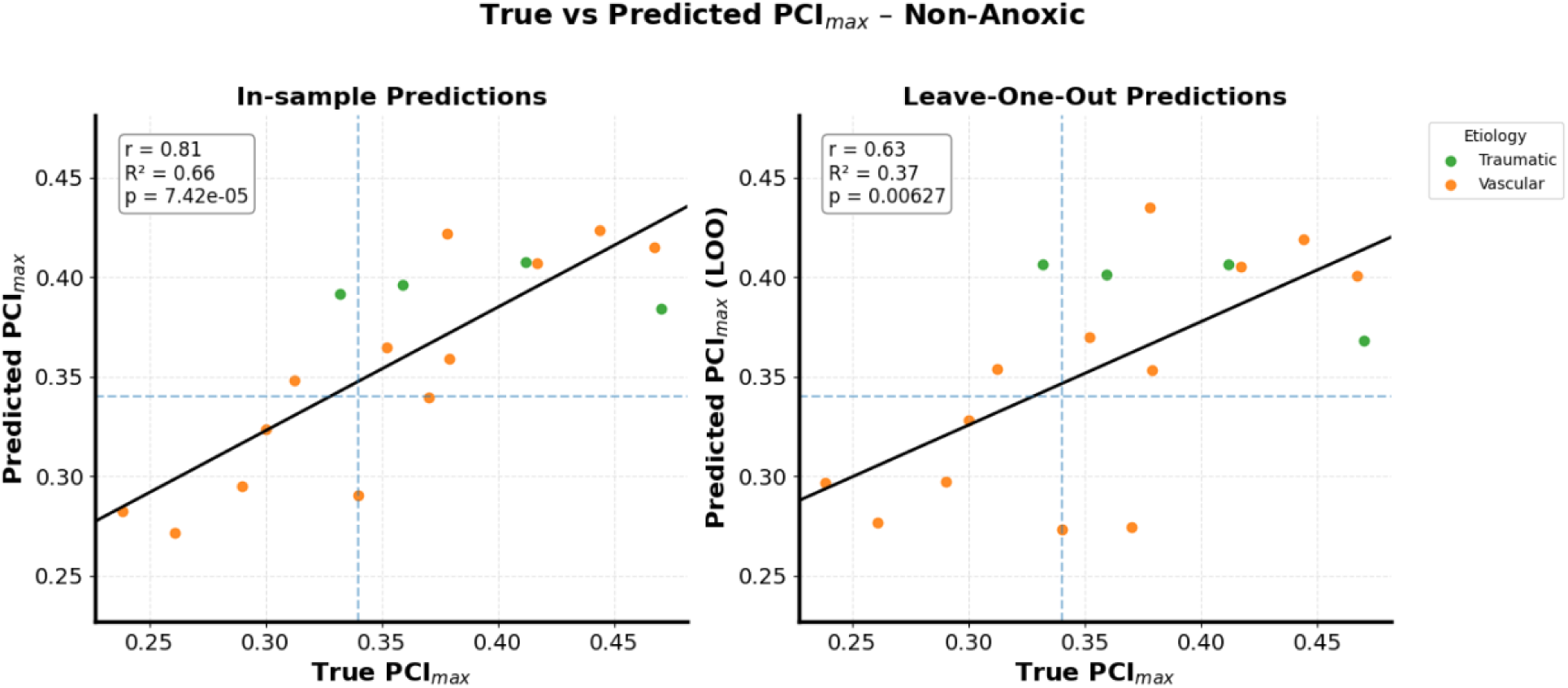
Baseline EEG criticality predicts perturbational complexity in non-anoxic patients. True versus predicted PCI_*max*_ values obtained from ridge regression models trained on resting-state EEG criticality features in non-anoxic patients (vascular and traumatic etiologies; n = 17). **Left**, in-sample predictions show a strong linear relationship between predicted and observed PCI_max_ values. **Right**, leave-one-out cross-validation (LOO-CV) predictions demonstrate preserved generalization, indicating that baseline criticality reliably predicts perturbational complexity across individuals. Points are colored by injury etiology (traumatic-green; vascular-orange). Dashed lines indicate the empirical PCI threshold (PCI = 0.34). These results show that when cortical architecture is relatively preserved, spontaneous critical dynamics constrain the system’s perturbational capacity.

Taken together, restricting the analysis to non-anoxic DoC etiologies reveals a stable and generalizable mapping of intrinsic dynamics of spontaneous EEG to the capacity to generate distributed, complex evoked responses to perturbation. This suggests that the failure observed in the full cohort are primarily driven by some anoxic patients with absent perturbational responses (PCI_max_ = 0), which introduces a pronounced floor effect that dominates prediction error loss (Fig. 2).

To determine whether the whole-cohort performance was driven primarily by floor effects rather than etiology, we performed a post-hoc analysis excluding patients with absent perturbational responses (PCI_*max*_= 0).

### Baseline criticality predicts PCI_*max*_ across etiologies when patients have a measurable perturbational response (PCI_*max*_ > 0)

In a post-hoc analysis, we tested whether predictive performance depended primarily on minimal preserved perturbational capacity, rather than etiology by excluding patients without measurable perturbational responses (PCI_*max*_ = 0). This yielded a responsive subgroup with non-zero PCI_*max*_ values (PCI_*max*_ > 0; *n* = 21) spanning all etiologies (vascular, *n* = 13; anoxic, *n* = 4; traumatic, *n* = 4). We therefore interpret PCI_*max*_ > 0 as the operational criterion to identify a subgroup with preserved measurable perturbational responsiveness.

Within this subgroup, the model again showed strong in-sample predictive performance (R² = 0.57; r = 0.758, p < 0.001). Although generalization was reduced relative to the non-anoxic group, LOO-CV performance remained statistically significant (R² = 0.07; MSE = 0.0059; MAE = 0.064; r = 0.44, p = 0.047) (Fig. 4). Recovering predictive performance across etiologies indicates that spontaneous EEG criticality may have predictive value for PCI_max_ when the cortex retains the capacity to generate a measurable perturbational response, even across heterogenous injury mechanisms.

**Figure 4.**
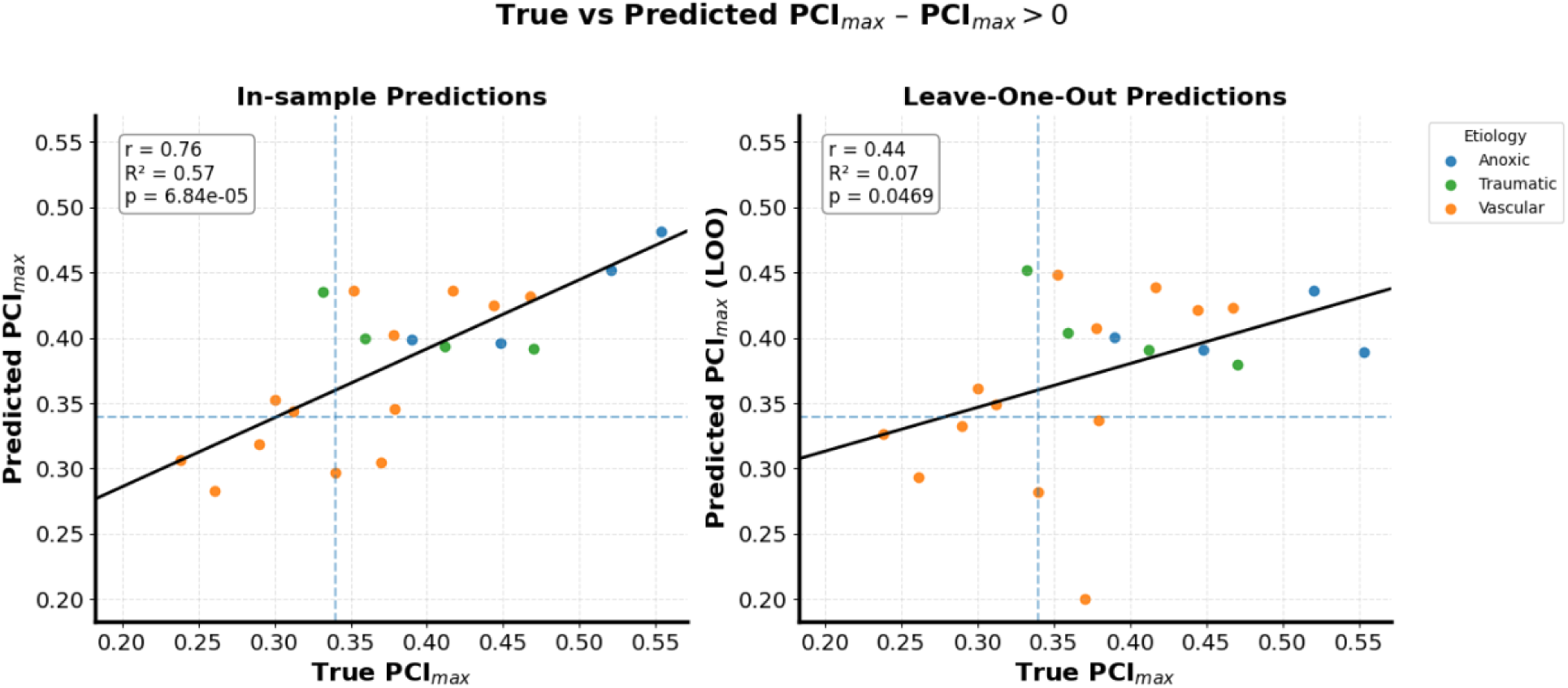
Baseline criticality predicts PCI_*max*_ in patients with measurable perturbational responses. True versus predicted PCI_*max*_ values for patients with non-zero PCI_*max*_ values (PCI_*max*_ > 0; n = 21), spanning all etiologies. **Left**, in-sample predictions reveal a strong association between baseline criticality features and PCI_*max*_. **Right**, LOO-CV predictions retain a significant but reduced correlation, indicating partial generalization. Points are colored by injury etiology (anoxic-blue; traumatic-green; vascular-orange). Dashed lines denote the PCI threshold (PCI = 0.34). These results indicate that spontaneous EEG criticality predicts perturbational complexity whenever the cortex retains structural integrity to generate a measurable response.

Together, these findings suggest that PCI_*max*_can be captured by a reproducible multivariate relationship within spontaneous criticality feature space, but that this mapping is strongly biased by PCI_*max*_ = 0 floor effects that compress the target range and destabilize individual prediction.

### Interdependent relation between criticality features

To characterize relationships among criticality features—the structure within the criticality feature space, we computed pairwise correlations across all 26 patients and applied Bonferroni correction to control for multiple comparisons (Fig. 5). The resulting matrix revealed highly structured interdependencies, indicating that EEG criticality is expressed across families of measures rather than isolated features.

**Figure 5.**
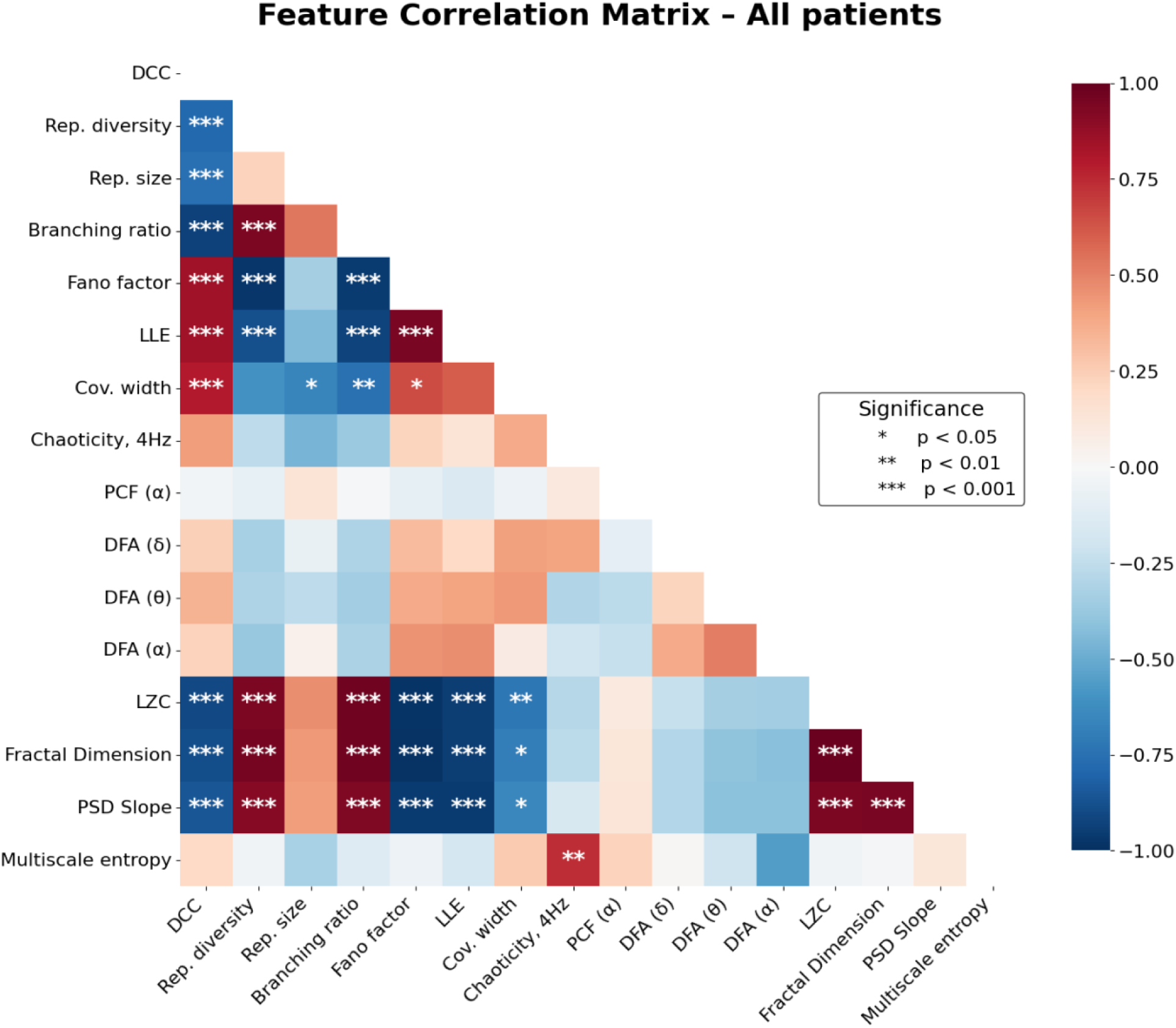
Criticality features exhibit structured collinearity across the full patient cohort. Pairwise Pearson correlation matrix computed across all EEG criticality features in the full cohort (n = 26). The matrix reveals heterogenous pattern of associations, with subsets of features showing strong positive or negative correlations, while others remain weakly related. Measures related to signal complexity and fractality (e.g., Lempel–Ziv complexity, fractal dimension, PSD slope) tend to covary, as do features associated with avalanche criticality (e.g., branching ratio, Fano factor, repertoire metrics), whereas temporal-scaling measures (e.g., DFA exponents, PCF α) show weak relationships. Color indicates correlation strength, with statistical significance denoted by asterisks. Together, these patterns indicate that EEG criticality is expressed through a coordinated but not redundant dynamical feature set rather than a single unified metric.

We observed families of covarying features: first, avalanche criticality measures—including deviation from criticality coefficient (DCC), branching ratio, Fano factor, repertoire size/diversity reflecting shared sensitivity to local propagation and event statistics; second, scaling and complexity measures—including Lempel–Ziv complexity, fractal dimension, and spectral slope (but not multiscale entropy), reflecting scale-free structure and compressibility in spontaneous temporal dynamics. Each of these families of features have strong interdependent positive correlations (|r| ≈ 0.6–0.9). In contrast, the measures relating to edge-of-synchrony and temporal scaling (Pair correlation function [PCF-alpha] and detrended fluctuation analysis [DFA - delta, alpha, and theta]) linked to oscillatory patterns and long-range temporal organization showed weak coupling with other features.

These interdependency patterns are consistent with the feature organization reported by Maschke et al.,^14^ and support the view that EEG criticality occupies a coordinated multivariate manifold rather than a single feature axis. Importantly, similar subgroup-specific correlation matrices were observed in the non-anoxic and PCI_max_ > 0 cohorts (Supplementary Fig. 1), indicating that the internal organization of the feature space is largely preserved across subgroups. Thus, the failure of whole-cohort prediction is unlikely to reflect a loss of structure within the criticality manifold itself, but rather a breakdown in how this structured spontaneous feature space maps onto PCI_max_ across response regimes.

### Univariate criticality–PCI_*max*_ correlations strengthen and converge in responsive subgroups

To further interpret subgroup prediction differences, we examined univariate associations between spontaneous EEG criticality features and PCI_*max*_ across the full cohort and restricted subgroups, where we performed Bonferroni correction for multiple comparisons per subgroup (Fig. 6; Supplementary Figs. 2-4). In the full cohort, individual feature-PCI_*max*_ relationships were weak and variable, and none of the 16 features survived Bonferroni correction, consistent with marked between-subgroup heterogeneity and floor compression at the lower bound of PCI_max_.

**Figure 6.**
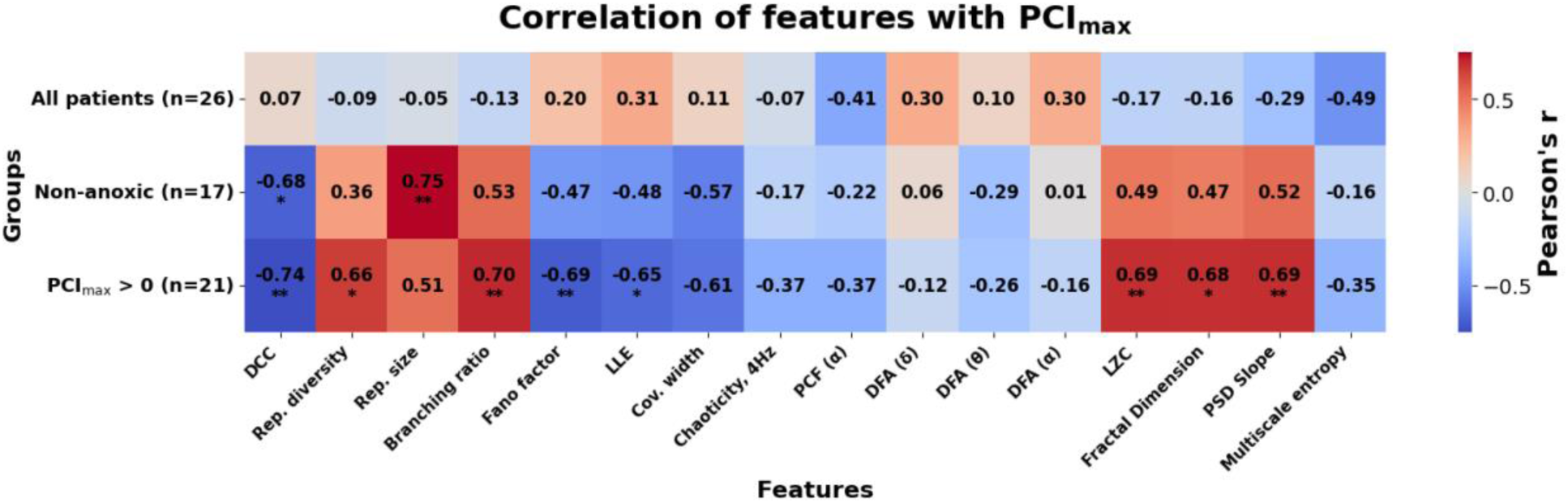
Complementary criticality features correlate to PCI_*max*_ in a subgroup-dependent manner. Heatmap showing Pearson correlations between individual EEG criticality features and PCI_*max*_ across three groups: all patients, non-anoxic patients, and patients with PCI_*max*_ > 0. While correlations are weak and inconsistent when all etiologies are pooled, multiple features show strong associations with PCI_*max*_ in non-anoxic and PCI_*max*_ > 0 subgroups. These patterns mirror the multivariate prediction results and indicate that perturbational complexity is linked to a constellation of complementary criticality features, contingent on preserved cortical structure.

In the non-anoxic subgroup, the relationships became more coherent and approximately linear, with 2 of 16 features remaining significant after correction: DCC (*r* = −0.68, *p*_Bonf_ = 0.046) and repertoire size (*r* = 0.75, *p*_Bonf_ = 0.03). Excluding all participants with PCI_*max*_ = 0 revealed substantially stronger and more consistent linear associations, with 8 of 16 features surviving Bonferroni correction, including DCC, repertoire diversity, branching ratio, Fano factor, LLE, LZC, fractal dimension, and PSD slope (all *p*_Bonf_ < 0.05). These findings mirror the partial recovery of predictive performance in multivariate regression and indicate that floor effects at PCI_max_ = 0, rather than etiology alone obscure an otherwise coherent relationship between spontaneous criticality dynamics and perturbational complexity.

Notably, the strongest single-feature correlations in each subgroup (all-patients: r = -0.49, non-anoxic: r = 0.75; PCI_*max*_ > 0: r = -0.74) did not exceed the in-sample performance of the multivariate ridge model (all-patients: r = 0.6, non-anoxic: r = 0.81, PCI_*max*_ > 0: r = 0.76) (Fig. 3 and 4), indicating that PCI_*max*_ is best captured by a joint multifeature relationship rather than any individual feature alone.

Together, these analyses demonstrate that spontaneous EEG criticality features become increasingly informative of perturbational complexity when cortical architecture is sufficiently preserved to elicit a measurable evoked response. In such cases, resting-state dynamics provide a reliable window into the brain’s perturbational capacity through TMS-EEG propagation, linking intrinsic dynamical regimes to externally evoked dynamical response.

### Mapping of behavioral and functional outcomes onto latent criticality manifold

As a post-hoc exploratory analysis, we next examined whether intrinsic critical dynamics related to clinically meaningful measures of consciousness and recovery. Behavioral responsiveness was assessed using the Coma Recovery Scale-Revised (CRS-R), where higher scores indicate greater behavioral responsiveness, and functional outcome was assessed using the Glasgow Outcome Scale-Extended (GOS-E), where higher scores indicate better functional outcome. All participants had CRS-R scores, but four participants lacked GOS-E scores, yielding a reduced analysis set of *n* = 22 for latent-space analyses involving functional outcome.

To characterize the geometry underlying the criticality feature space, we embedded participants into a low-dimensional latent manifold using principal component analysis (PCA) on the standardized 16 feature matrix (Fig. 7A-C). The first three components explained 80.1% of total variance (PCA-1 = 55.4%, PCA-2 = 16.5%, PCA-3 = 8.2%), indicating that spontaneous criticality features occupy a compact latent manifold rather than spanning independent axes. Feature loadings indicate that PCA-1 captures a global criticality organization dominated by avalanche criticality measures (e.g., repertoire diversity/size, branching ratio), alongside complexity and fractal scaling (e.g., Lempel–Ziv complexity, fractal dimension), whereas PCA-2 captures an orthogonal axis with opposing loading structure across several features (Fig. 7C). Subjects were distributed continuously across this manifold, with partial separation by etiology along the dominant axes (Fig. 7A).

**Figure 7.**
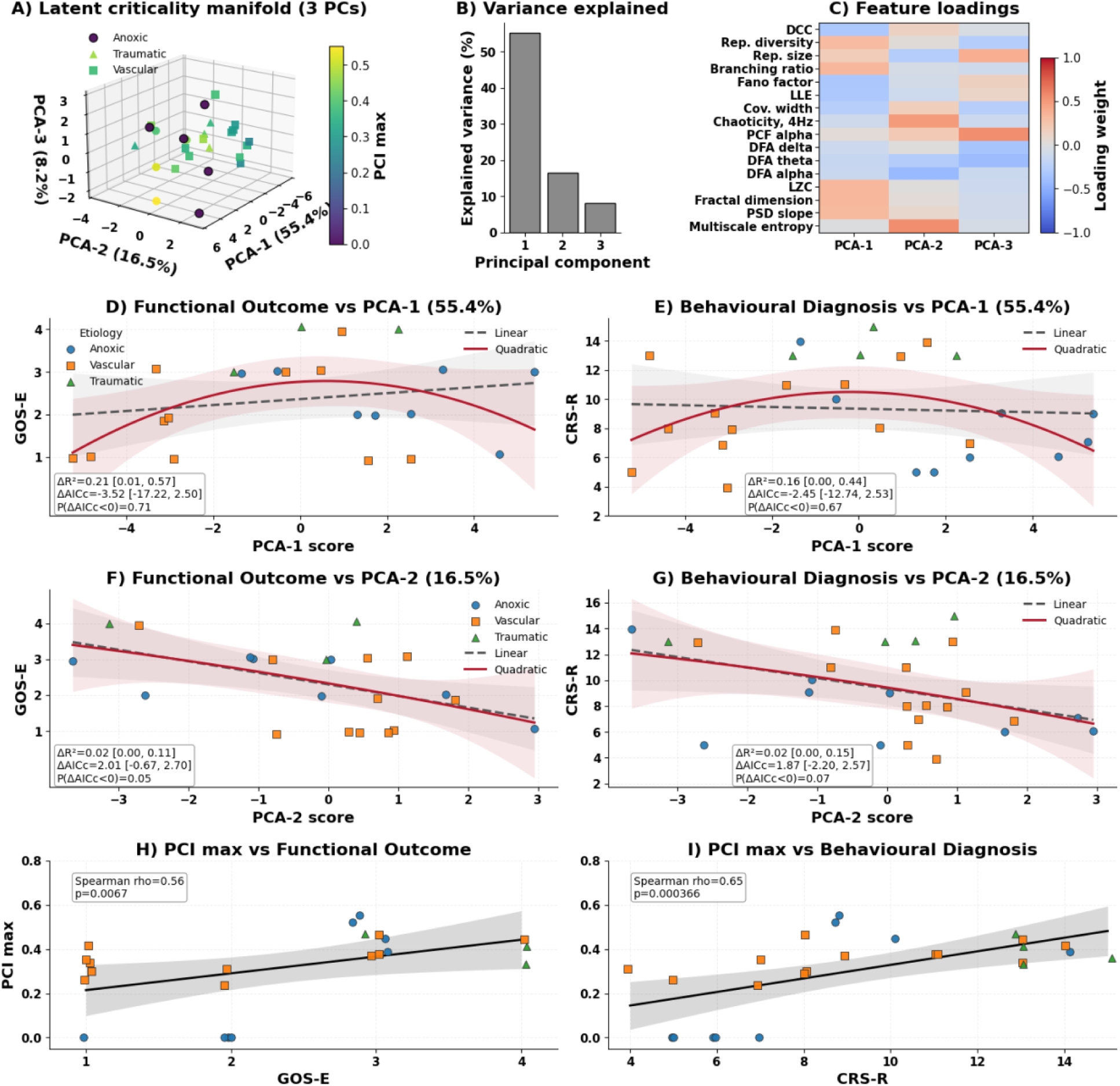
Latent criticality manifold of spontaneous EEG dynamics and its relationships with behavioural responsiveness, functional outcome and PCI_*max*_. (A) Three-dimensional principal component analysis (PCA) of 16 resting-state EEG criticality features, with each point representing one patient and colored by PCI_*max*_. Marker colour denotes etiology (anoxic, vascular, traumatic). The first three principal components define a latent criticality manifold summarizing coordinated variation across features. (B) Variance explained by the first three principal components. (C) Loadings of individual EEG criticality features onto PCA-1–PCA-3, showing the multivariate feature structure underlying the latent space. (D–G) Associations between latent criticality axes and clinical measures. PCA-1 was related to functional outcome measured by GOS-E (D) and behavioral responsiveness measured by CRS-R (E), higher values on both scales indicate better clinical status. PCA-2 was related to GOS-E (F) and CRS-R (G). For each PCA-clinical association, ordinary least squares models were fit using both linear and quadratic forms. Grey dashed lines show the linear fit, red solid lines show the quadratic fit, and shaded bands indicate 95% confidence intervals estimated by bootstrap resampling of the fitted curves. Text boxes report bootstrap model-comparison statistics: the mean change in explained variance 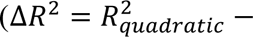 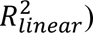, the mean change in small-sample corrected Akaike information criterion (ΔAICc = *AICc*_*quadratic*_ − *AICc*_*linear*_), and the proportion of bootstrap resamples in which the quadratic model was preferred (P(ΔAICc < 0)). Bracketed intervals indicate empirical 95% bootstrap intervals, calculated from the 2.5^th^ and 97.5^th^ percentiles of the bootstrap distribution. Negative ΔAICc values support the quadratic model relative to the linear model. Points are coloured and shaped by etiology. (H–I) PCI_max_ was positively associated with functional outcome (GOSE; H) and behavioral responsiveness (CRS-R; I). Small horizontal jitter was applied to GOS-E and CRS-R values for visualization only, to reduce overlap among points sharing the same score. Black lines indicate trend fits, gray shading denotes confidence intervals, with reported Spearman correlation coefficients and p-values. Together, these results suggest that baseline EEG criticality is expressed as a low-dimensional latent manifold that relates to both spontaneous behavioral state and the brain’s capacity for complex perturbational responses.

We then mapped behavioural responsiveness and functional outcome onto these latent axes (Fig. 7D-G). Along PCA-1, both GOS-E and CRS-R showed predominantly nonlinear relationships. For GOS-E, the linear model explained little variance (*R*^2^ = 0.04), whereas inclusion of a quadratic term improved the fit (*R*^2^ = 0.22; Δ*R*^2^ = 0.18). For CRS-R, the linear model explained little variance (*R*^2^ = 0.03), whereas the quadratic model explained more variance (*R*^2^ = 0.14; Δ*R*^2^ = 0.11). Bootstrap resampling supported this improvement for both outcomes (GOS-E: mean Δ*R*^2^ = 0.21 [0.01, 0.57]; CRS-R: mean Δ*R*^2^ = 0.16 [0.00, 0.44]). The quadratic model was favoured by small-sample corrected Akaike information criterion (AICc) in 70.6% of bootstrap resamples for GOS-E (mean ΔAICc = −3.52 [−17.22, 2.5]) and 66.6% for CRS-R (mean ΔAICc = −2.45 [−12.74, 2.53]), suggesting moderate support for a nonlinear, inverted-U profile with better clinical status at intermediate PCA-1 values (Fig. 7D-E). Together, PCA-1 showed reproducible increases in explanatory fit with quadratic modeling, although AICc analysis showed moderate support as confidence intervals crossed zero.

In contrast, PCA-2 showed monotonic negative relationships with both clinical measures, with lower GOS-E (*R*^2^ = 0.27, *p* = 0.014; Fig. 7F) and lower CRS-R (*R*^2^ = 0.16, *p* = 0.0397; Fig. 7G). Adding a quadratic term provided negligible improvement (GOS-E: Δ*R*^2^ = 0.02 [0.00, 0.11], ΔAICc = 2.01 [−0.67, 2.7], *P*(ΔAICc < 0) = 0.05; CRS-R: Δ*R*^2^ = 0.02 [0.00, 0.15], ΔAICc = 1.87 [−2.2, 2.57], *P*(ΔAICc < 0) = 0.07). Unlike PCA-1, PCA-2 appeared to capture a more linear clinical association, where higher PCA-2 values were associated with lower functional outcome and lower behavioural responsiveness.

PCI_*max*_ showed strong associations with clinical measures (Spearman ρ = 0.56 with GOS-E, p = 0.0067; Spearman ρ = 0.65 with CRS-R, p < 0.001), consistent with PCI_*max*_ as an index of consciousness capacity (Fig. 7H-I).

Together, these analyses indicate that resting-state EEG criticality is organized as a low-dimensional dynamical manifold that maps onto clinically relevant outcome measures. PCA-1 captures a possible intermediate optimal regime, whereas PCA-2 captures more monotonic variation in behavioral and functional status. For completeness, associations between each individual EEG criticality feature and both GOS-E and CRS-R were examined (Supplementary Fig. 5-6).

## Discussion

In this study, we tested whether spontaneous EEG criticality features could predict perturbational complexity in patients with disorders of consciousness. We found that the relationship between spontaneous EEG criticality and perturbational complexity in DoC is conditional on cortical response regime. Spontaneous criticality features did not reliably predict PCI_*max*_ when all patients were pooled together, but prediction became robust in non-anoxic patients and partially recovered when analyses were restricted to patients with preserved perturbational responses (PCI_*max*_ > 0). These findings indicate that the relationship between spontaneous EEG criticality features and PCI_*max*_ is not a universal surrogate for PCI across DoC. Rather, spontaneous criticality reflects intrinsic dynamical organization that maps onto PCI_*max*_ only when the cortical architecture required to support propagation of an externally evoked perturbation is sufficiently preserved. When perturbational responses are absent and PCI_*max*_ collapses to a floor value, the spontaneous-evoked mapping becomes ill-posed.

Extending the findings from Maschke et al.,^14^ from anesthesia to DoC, these results help clarify how spontaneous and perturbational markers of consciousness are related. PCI_*max*_ indexes the brain’s capacity to generate a complex spatiotemporal response to external stimulation, whereas resting-state EEG criticality measures characterize the intrinsic dynamical regime in which the system operates. Therefore, PCI_*max*_ may be understood as an expression of the underlying intrinsic dynamical state capable of sustaining complex interactions. In this view, near-critical neural dynamics may enable the integration and differentiation thought to support consciousness^13,15,19^, while also providing the intrinsic conditions under which external perturbations can evoke complex cortical responses. In severe anoxic injury, spontaneous EEG criticality features remain measurable, but they no longer map onto PCI_*max*_ because the perturbational response is effectively saturated at the lower bound. This is consistent with previous findings that EEG recordings in anoxic patients reflect distinct pathophysiological mechanisms as revealed by suppressed alpha power^23^. More broadly, when assessing consciousness with PCI in patients with a DoC, the population should not be treated as a physiologically homogenous category and etiology should be considered^23,25,27^.

The univariate analyses further support our interpretation that spontaneous and perturbational indices are related, but not interchangeable. Individual features showed subgroup-dependent associations with PCI_*max*_, with stronger and more convergent correlations in non-anoxic and responsive subgroups than in the full cohort. However, the strongest single-feature associations did not exceed the performance of the multivariate ridge model, indicating that PCI_*max*_ is not captured by any single criticality feature alone. Rather, perturbational complexity appears to be best explained by a coordinated configuration of criticality features. This is consistent with the prior work suggesting that criticality in the brain is expressed as a multifeatured dynamical regime rather than a single observable^19,28–30^.

The multivariate feature structure provides an additional layer of interpretation. Overall, criticality features show coherent correlation patterns that partially resemble the healthy-anesthesia results in Maschke et al.,^14^ particularly the associations of feature families including avalanche criticality, edge-of-chaos, edge-of-synchrony and criticality-related features (Fig. 5). However, several differences were notable. Repertoire size shows limited significant correlations when anoxic patients are included, reflecting fragmented remnants of avalanche propagation in anoxic patients. Chaoticity is significantly correlated only with multiscale entropy, and DFA measured across the delta, theta, and alpha bands showed no significant relationships with the remaining feature set, suggesting a partial decoupling of edge-of-chaos and long-range temporal scaling features in severe brain injuries. Fano factor shows a systematic reversal in its correlations relative to healthy anesthesia, indicating that the variability-based measure may index a different dynamical regime in the injured cortex. Together, these findings suggest that severe brain injury weakens and partially reorganizes the latent structure linking avalanche, and edge-of-synchrony criticality measures.

The post-hoc diagnostic and prognostic analyses suggest that this latent structure may also be relevant beyond its correlation with PCI_*max*_. Behavioral responsiveness and long-term functional outcome appeared to map onto the intrinsic geometry of spontaneous EEG critical dynamics. Thus, the capacity for consciousness is unlikely to be maximized by a monotonic change in a single criticality feature but may instead depend on occupying an optimal operating regime along the dominant axis of criticality feature covariation. PCA-1 showed a relation with clinical variables that was better explained by an inverse quadratic, inverted-U pattern. Bootstrap resampling showed that quadratic models captured more variance for both GOS-E and CRS-R. However, AICc support was moderate and this nonlinear pattern should be interpreted cautiously. Conceptually, the inverted-U profile suggests that behavioural responsiveness and functional recovery are greatest within an intermediate dynamical regime balanced between excessive order or disorder. In this view, coordinated shifts across criticality features may reflect neural activity between overly constrained, subcritical-like dynamics, and overly unstable, supercritical-like dynamics, with an intermediate regime facilitating flexible brain responses that support consciousness^15,19^ (Fig. 7).

Although exploratory, this result suggests that clinically meaningful variation may reflect regimes within a broader dynamical landscape in severe brain injury. The anoxic subgroup was notably separable in both clinical variables to the positive domain of PCA-1. In contrast, the relationships between PCA-2 and both CRS-R and GOS-E were better explained by linear models, reflecting a more monotonic dimension of pathological organization associated with worse behavioural measures of consciousness and functional recovery. Nevertheless, PCI_*max*_ had stronger associations with clinical variables than either latent axis alone in this cohort. However, PCI has an important limitation: it depends on the ability to elicit, propagate, and measure a cortical response to perturbation. Consequently, a floor-level PCI response indicates the absence of a measurable perturbational response but does not necessarily indicate an absence of residual network capacity or potential recovery of consciousness. Such floor-level values may arise from multiple mechanisms, including reduced cortical excitability, low signal-to-noise ratio, injury-related suppression, or technical limitations in detecting the response.

Spontaneous critical dynamics may provide complementary information that reveal latent capacity for recovery of consciousness when perturbational measures are limited by floor effects. Practically, PCI is one of the most informative available markers of consciousness, but its acquisition requires TMS-EEG and specialized expertise that limit bedside scalability. Resting-state EEG is far easier to acquire in clinical settings. Our results therefore suggest that passive EEG criticality may offer a tractable marker of perturbational capacity. In clinical practice, resting-state EEG could serve as a prior to guide TMS-EEG and complement the PCI. At the same time, the failure of whole-cohort prediction cautions against assuming that a single passive EEG model will generalize across all forms of severe brain injury. Response regime, etiology and structural integrity likely need to be accounted for explicitly when considering passive spontaneous EEG models of PCI.

Several limitations should be acknowledged. First, the sample size was modest, especially after stratification by etiology and perturbational responsiveness. Second, the subgroup restriction to PCI_*max*_ > 0 was post hoc and should therefore be interpreted as exploratory rather than confirmatory. Third, the present analyses relied on linear ridge regression, which may underestimate nonlinear relationships between spontaneous criticality features and perturbational complexity. This may be especially relevant in heterogeneous etiologies, where floor effects, saturation, and regime shifts could distort linear mappings, and future studies should consider non-linear methods in larger cohorts. Fourth, the regularization hyperparameter was set to α = 1, a moderate default regularization strength, future studies should optimize this value and determine the optimal predictive performance on a larger sample. Finally, the use of PCA to construct the latent criticality axes may be influenced by the redundancy among correlated features. Future work should test whether similar latent criticality structure and clinical associations emerge using approaches such as factor analysis or feature clustering.

In summary, resting-state EEG criticality predicted perturbational complexity only under conditions in which cortical responsiveness was sufficiently preserved to elicit a PCI measurement. Prediction failed in the pooled cohort largely because floor-level PCI_max_ values disrupted the mapping between intrinsic spontaneous dynamics and externally evoked complexity. Anoxic injury provided a clinically important example of this suppressed regime, but the relevant mechanistic distinction was preserved perturbational propagation. More broadly, they support the idea that passive EEG may provide a scalable window into the brain’s capacity for complex, consciousness-supporting dynamics.

## Methods

### Participants and clinical assessment

Data were obtained from a previously published multicenter dataset of patients in disorders of consciousness (DOC), collected across several clinical sites and approved by local institutional ethics committees^4,23^. Consent was obtained from legal representatives. We analyzed a subset of 26 patients from this dataset (20 females and 6 males, 22-76 years old) who had complete resting-state EEG recordings and available TMS-EEG derived Perturbational Complexity index (PCI) estimates.

Patients were included at least one week post-injury and spanned acute and chronic DOC, with a range of 0.43-128.94 months since injury (mean = 25.63 months). Clinical behavioral responsiveness scores were determined from repeated administration of the Coma Recovery Scale-Revised (CRS-R), with the final score based on the best observed behavioural response across multiple assessments within the same evaluation week. EEG recordings were acquired at least 20 days after injury onset and at least 3 days after sedation withdrawal^25^. Functional outcome measurements were assessed at 6 months post-injury with the Glasgow Outcome Score – Extended (GOS-E) and only available for 22 patients^26^. The cohort comprised patients with three primary etiologies, vascular injuries were the most common (*n* = 13), followed by anoxic injuries (*n* = 9) and traumatic injuries (*n* = 4). All patients were free from sedative medication at least seven days prior to EEG acquisition.

### Electroencephalography data

#### Acquisition

Resting state EEG was recorded while patients were in a clinical environment. Spontaneous EEG recordings (mean duration = 6.45 min, interquartile range = 4.26-8.02 min) were acquired on the same day as the TMS-EEG protocol or within a time window where behavioural assessments remained stable across session. EEG data were recorded using TMS-compatible amplifiers (Nexstim Ltd. (*n* = 23) or Brain Products GmbH (*n* = 3)) with 60 or 64 scalp electrodes arranged in comparable montages and sampling rates of 1,450 Hz or 5,000 Hz, respectively. During acquisition electrode impedance was maintained below 20kΩ with a reference near Fpz.

#### Preprocessing

EEG data were preprocessed prior to the present analysis following the pipeline described in Colombo et al.,^23^. Briefly, EEG signals were band-pass filtered using a zero-phase Butterworth filter (0.5–60 Hz) and notch-filtered to attenuate line noise (50 Hz) and its harmonics. Non-physiological segments were identified through visual inspection by trained experts and removed. Channels exhibiting persistent noise were excluded and interpolated with spherical spline. All recordings were then average referenced. To further reject residual artifacts, independent component analysis (ICA) was performed, and components corresponding to ocular, muscular, or cardiac activity were identified and removed prior to signal reconstruction. Additional preprocessing step included downsampling to 500 Hz and segmentation into non-overlapping 10-second epochs. These steps ensured high signal fidelity while preserving broadband neural dynamics relevant for criticality analysis.

#### EEG feature extraction

From each preprocessed resting-state EEG recording, we estimated a set of 16 features designed to quantify aspects of neural criticality and dynamical organization. For each recording up to the first five minutes were used for feature extraction. Features were computed at the channel and epoch level and then averaged to obtain a single subject-level value. These features span avalanche criticality (deviation from criticality coefficient (DCC), repertoire size and diversity, branching ratio, and Fano factor), edge of chaos criticality (chaoticity estimates, largest Lyapunov exponent (LLE), and width of covariance matrix), and criticality-related measures (spectral slope, fractal dimension, Lempel–Ziv complexity, multiscale entropy, detrended fluctuation analysis (DFA) (applied to the alpha, delta and theta band-passes) and pair correlation function (PCF) in the alpha band. Features were computed using the same Python-based analysis pipelines introduced in Maschke et al.,^14^. For a complete description of each feature see Maschke et al.,^14^.

### Perturbational Complexity Index

Perturbational Complexity Index (PCI) values were obtained from prior publications^4,23^. PCI quantifies the algorithmic complexity of spatiotemporal cortical responses to external perturbation via transcranial magnetic stimulation (TMS). TMS-evoked dynamical responses are thresholded to isolate and binarize significant cortical responses. The resulting binary matrix is then compressed using the Lempel-Ziv algorithm normalized by source entropy, see Casali et al.,^2^ for a detailed description of the protocol. The maximum PCI value across stimulation sites (PCI_*max*_) was used in the analysis. For subgroup analyses, PCI_*max*_ > 0 was used as an operational marker of a measurable perturbational response, rather than as a characterization of the physiological quality or the spatial propagation of the TMS-EEG response.

### Predictive modeling and statistical analysis

We evaluated whether spontaneous EEG criticality features could predict PCI_*max*_ using multivariate ridge regression. The model inputs included the full 16-feature set, with PCI_*max*_ as a continuous prediction target. The regularization parameter was fixed at α = 1 across all analyses.

Model performance was evaluated using both in-sample prediction and leave-one-out cross-validation (LOO-CV). In-sample fits were used to characterize learned relationship between resting-state features and PCI_*max*_, while LOO-CV was used to assess generalization to unseen subjects. Predictive performance was quantified using the coefficient of determination (R²), mean squared error (MSE), mean absolute error (MAE), and Pearson correlation between predicted and observed PCI_*max*_ values.

To assess the impact of injury heterogeneity, models were evaluated separately across three groups: the full cohort (*n* = 26), non-anoxic patients only (*n* = 17), and patients with non-zero PCI_*max*_ values (*n* = 21). This stratified analysis was designed to determine whether preserved cortical responsiveness and etiology influenced the ability of spontaneous EEG dynamics to predict perturbational capacity.

### Correlation structure among features

To characterize the internal dependency structure of the criticality feature space, we computed pairwise Pearson correlation coefficient between all 16 subject-level EEG criticality features across the full cohort. Statistical significance was assessed using Bonferroni correction. The same analysis was repeated within the non-anoxic and PCI_*max*_ > 0 subgroups to determine whether the internal organization of the feature space was preserved across stratified cohorts.

### Univariate analyses

Additionally, pairwise Pearson correlations were computed between individual EEG features and PCI_*max*_ to assess univariate associations. Statistical significance was evaluated using Bonferroni correction to control the family-wise error rate for each tested feature-feature and feature-clinical measure pairs.

### Latent criticality manifold and post-hoc clinical association analyses

To characterize the low-dimensional organization of the subject-level EEG criticality feature space, we performed principal component analysis (PCA) after standardizing each feature across subjects to zero mean and unit variance. For visualization of the latent criticality manifold, PCA was applied to the full cohort feature matrix, and the first three principal components were retained for three-dimensional representation.

For post-hoc analyses relating latent criticality structure to clinical measures, we restricted the dataset to participants with available functional outcome data (n = 22) and PCA was recomputed within this reduced cohort. Subject scores on PCA-1 and PCA-2 were then related to functional outcome (GOS-E) and behavioral responsiveness (CRS-R).

Associations involving PCA-1 were modeled using ordinary least squares (OLS) linear and quadratic regression. The linear model was specified as *y* = β_0_ + β_1_*x*, and the quadratic model as *y* = β_0_ + β_1_*x* + β_2_*x*^2^, where *x* denotes the principal component score and *y* the clinical measure. Model comparison was summarized using the coefficient of determination (*R*^2^) and the change in explained variance between linear and quadratic models 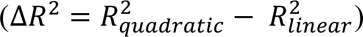. To evaluate whether the additional quadratic term improved model fit considering increased model complexity, we compared linear and quadratic models using the small-sample corrected Akaike Information Criterion (AICc) (ΔAICc = *AICc*_*quadratic*_ − *AICc*_*linear*_). Where, negative ΔAICc values indicate support for the quadratic model. The stability of model improvement was assessed using nonparametric bootstrap resampling. For each PCA-clinical measure association, paired predictor-outcome observations were resampled with replacement 1,000 times. Within each bootstrap sample, both linear and quadratic models were refit, and bootstrap distributions for Δ*R*^2^ and ΔAICc were obtained. For both quantities, pointwise 95% confidence intervals were obtained from the empirical 2.5th and 97.5th percentiles of the corresponding bootstrap distribution values. The proportion of bootstrap samples with ΔAICc < 0 was used to quantify how consistently the quadratic model was preferred over the linear model.

To assess direct relationships between perturbational complexity and behavioural measures, PCI_max_ was related separately to GOS-E and CRS-R, associations were summarized using Spearman rank correlations. Additionally, direct relationships between each resting-state EEG criticality feature and behavioural measures were assessed using spearman rank correlations.

