## Supplementary Materials for "Critical dynamics in spontaneous EEG predict perturbational complexity in disorders of consciousness with measurable evoked responses"

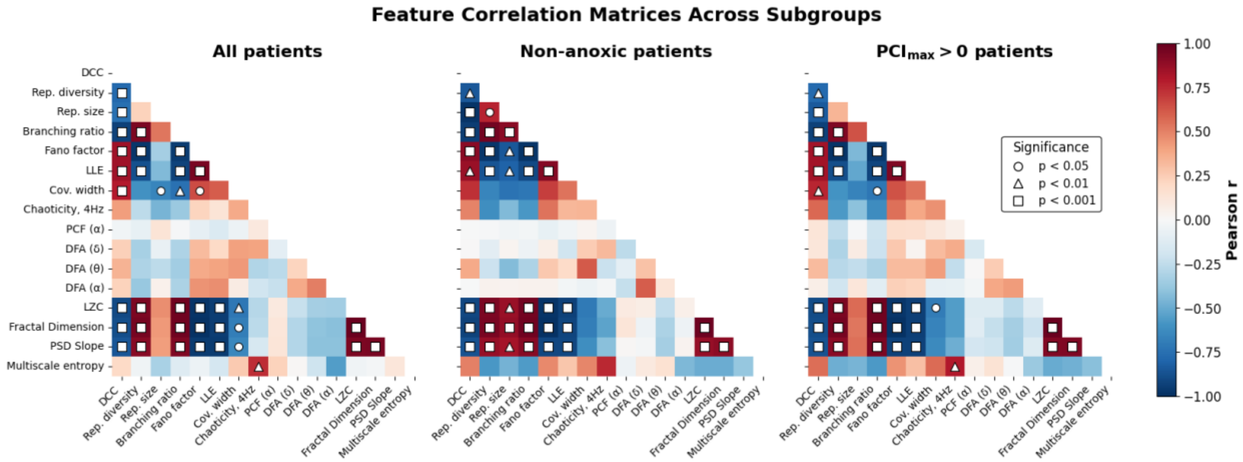


**Supplementary Figure 1. Criticality features exhibit structured collinearity across all subgroups.** Pairwise Pearson correlation matrix computed across all EEG criticality features in the full cohort ($n$ = 26), the non-anoxic subgroup ($n$ = 17), and the $\mathrm{PCI}_{\mathrm{max}} > 0$ subgroup ($n$ = 21). All subgroup matrices reveal heterogenous pattern of associations, with subsets of features showing strong positive or negative correlations, while others remain weakly related. Measures related to signal complexity and scaling structure, including Lempel-Ziv complexity, fractal dimension, and PSD slope, tended to covary, as did several features associated with avalanche criticality, including branching ratio, Fano factor, and repertoire metrics. By contrast, temporal-scaling and synchrony-related measures, such as DFA exponents and PCF-$\alpha$, showed weaker and less consistent relationships with the rest of the feature space. Color indicates correlation strength ($r$), and symbol overlays denote Bonferroni-corrected significance levels. Overall, these matrices indicate that EEG criticality is expressed through a coordinated but non-redundant multivariate feature set rather than a single unified metric, and that this internal structure is preserved across subgroups despite differences in their relationship to $\mathrm{PCI}_{\mathrm{max}}$.

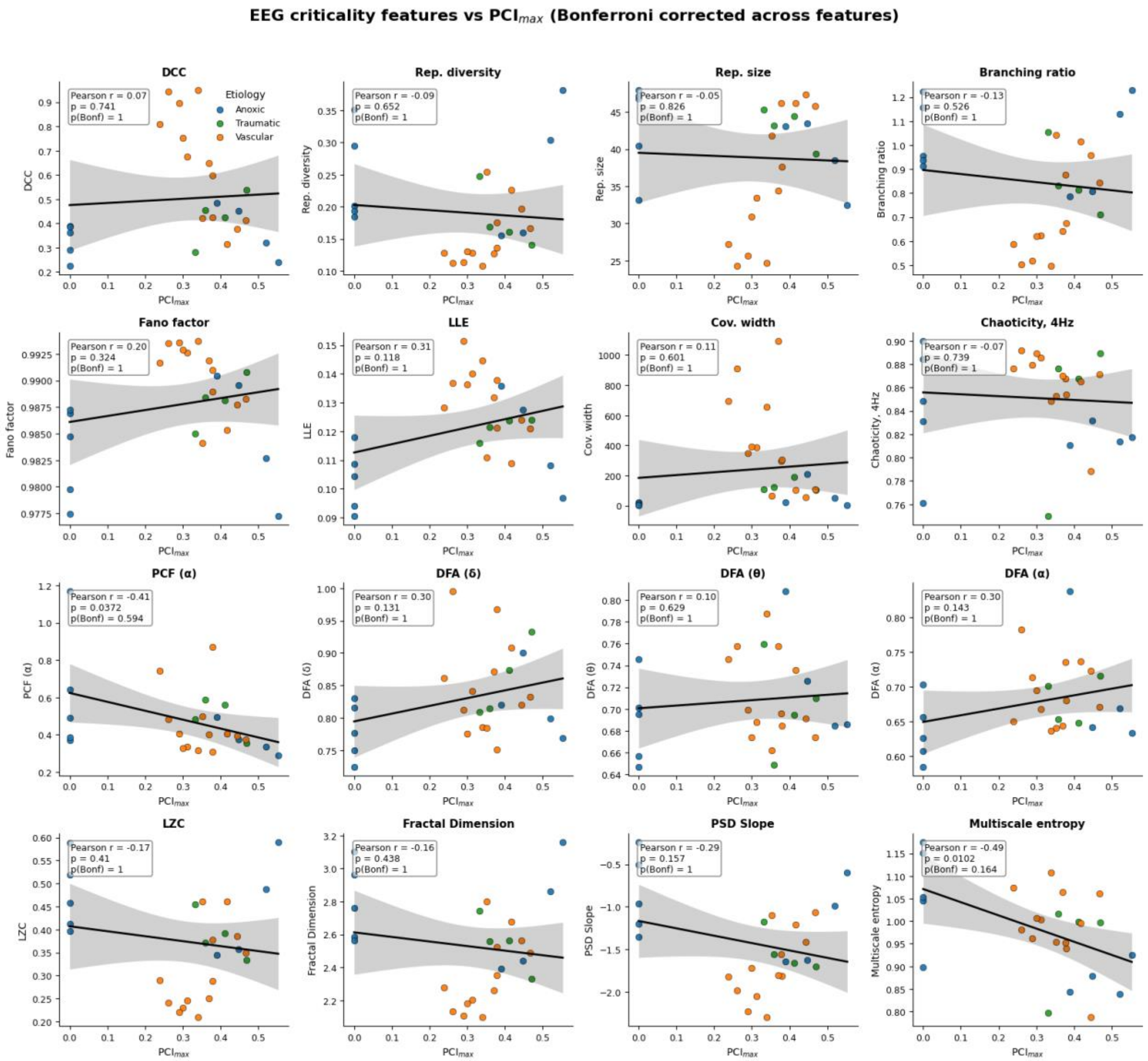


**Supplementary Figure 2. Spontaneous EEG criticality features and PCI$_{max}$ in the full cohort.** Each panel shows the relationship between PCI$_{max}$ and one EEG criticality feature. Points are colored by etiology (anoxic, vascular, traumatic), the solid black line shows the least-squares linear fit, and the shaded band indicates the 95% confidence interval of the fitted mean. Reported Pearson's correlation coefficient ($r$), the nominal $p$-value, and the Bonferroni-corrected $p$-value computed for all features in the grid.

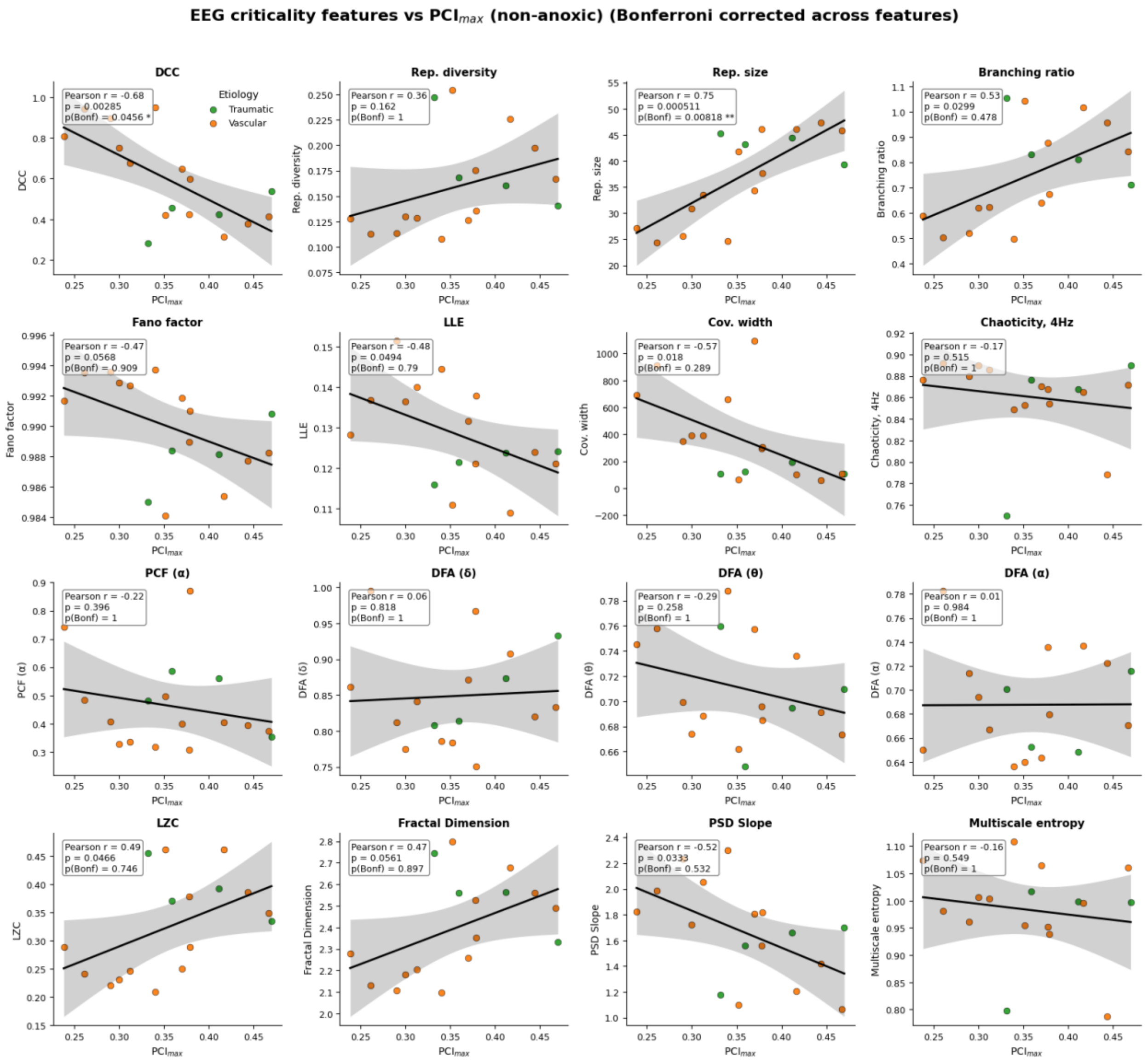


**Supplementary Figure 3. Spontaneous EEG criticality features and PCI$_{max}$ in the non-anoxic subgroup.** Each panel shows the association between one spontaneous EEG criticality feature and PCI$_{max}$ after excluding anoxic participants. Points are colored by etiology (anoxic, vascular, traumatic), the solid black line shows the least-squares linear fit, and the shaded band indicates the 95% confidence interval of the fitted mean. Reported Pearson's correlation coefficient ($r$), the nominal $p$-value, and the Bonferroni-corrected $p$-value computed for all features in the grid.

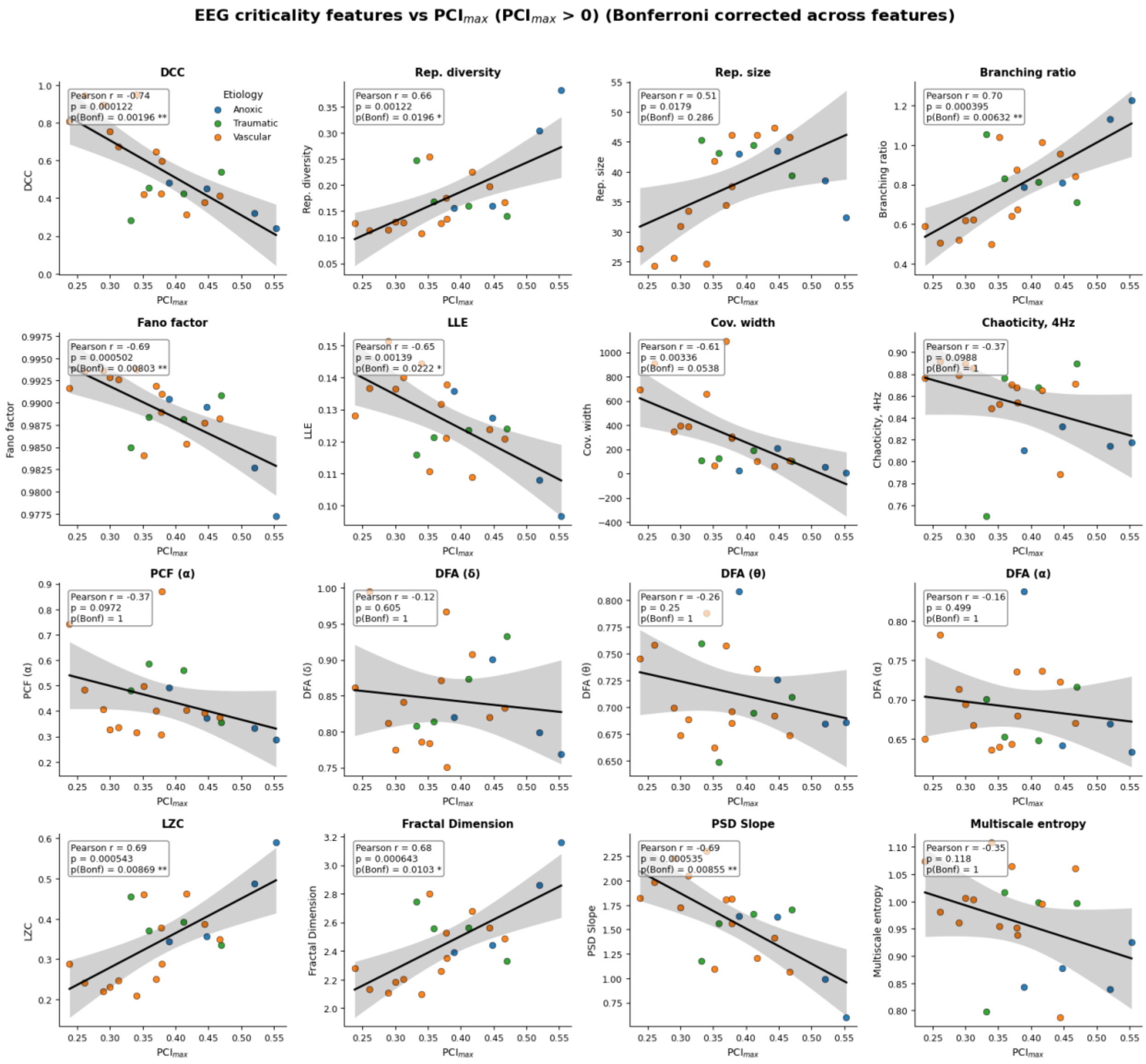


**Supplementary Figure 4. Spontaneous EEG criticality features and $PCI_{max}$ in participants with non-zero perturbational responses ($PCI_{max} > 0$).** Each panel shows the association between one spontaneous EEG criticality feature and $PCI_{max}$ after excluding participants with $PCI_{max} = 0$. Points are colored by etiology, black lines indicate the least-squares linear fit, and gray shading denotes the 95% confidence interval. Reported Pearson's correlation coefficient ($r$), the nominal $p$-value, and the Bonferroni-corrected $p$-value computed for all features in the grid.

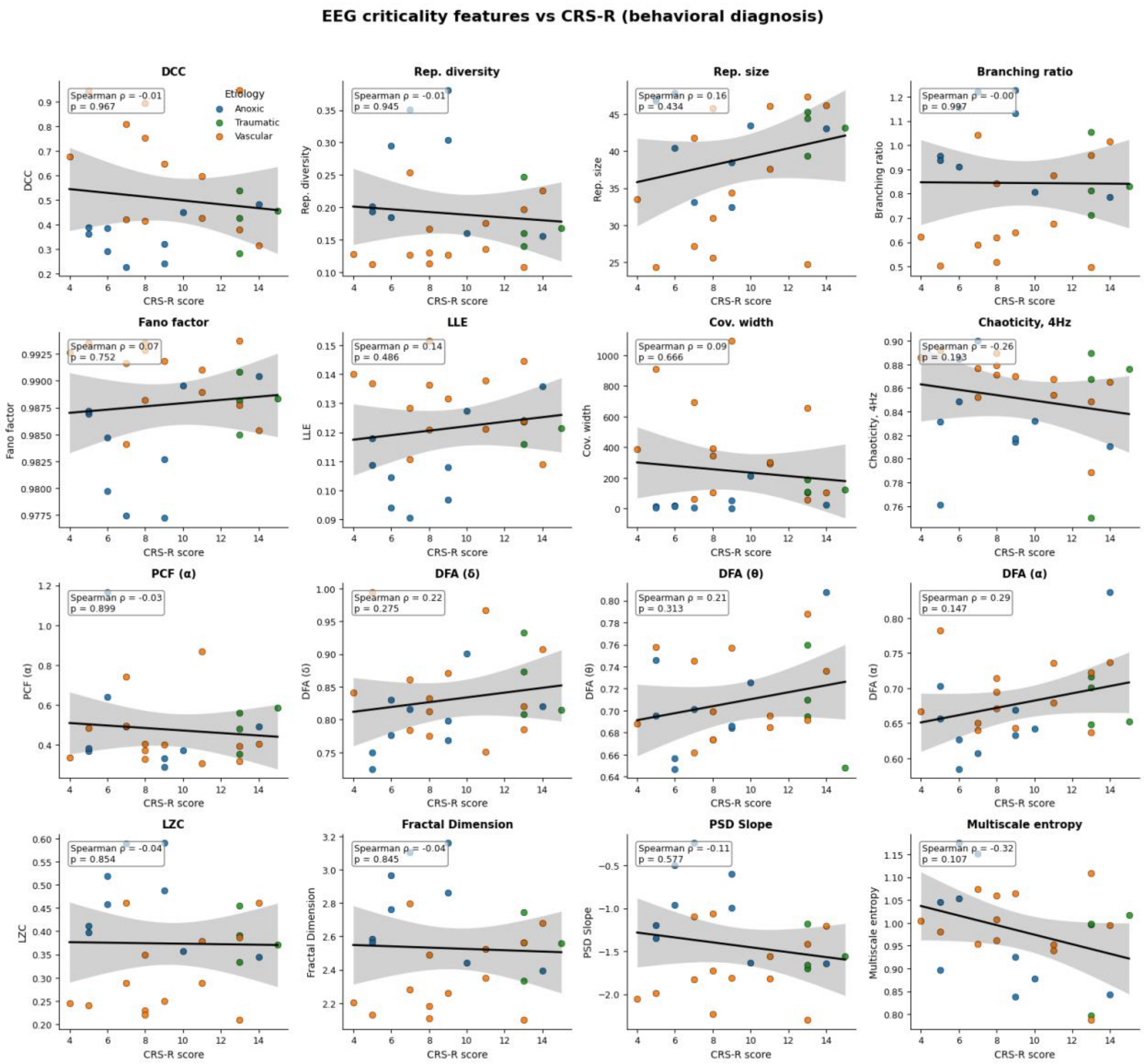


**Supplementary Figure 5. Associations between individual EEG criticality features and behavioral responsiveness (CRS-R).** Each panel shows the relationship between one resting-state EEG criticality feature and behavioral responsiveness as measured by the Coma Recovery Scale–Revised (CRS-R). Points represent individual patients and are colored by etiology (anoxic, vascular, traumatic). Black lines indicate fitted linear trend lines and gray shading denotes 95% confidence intervals. Each plot reports Spearman rank correlation coefficients and associated p-values.

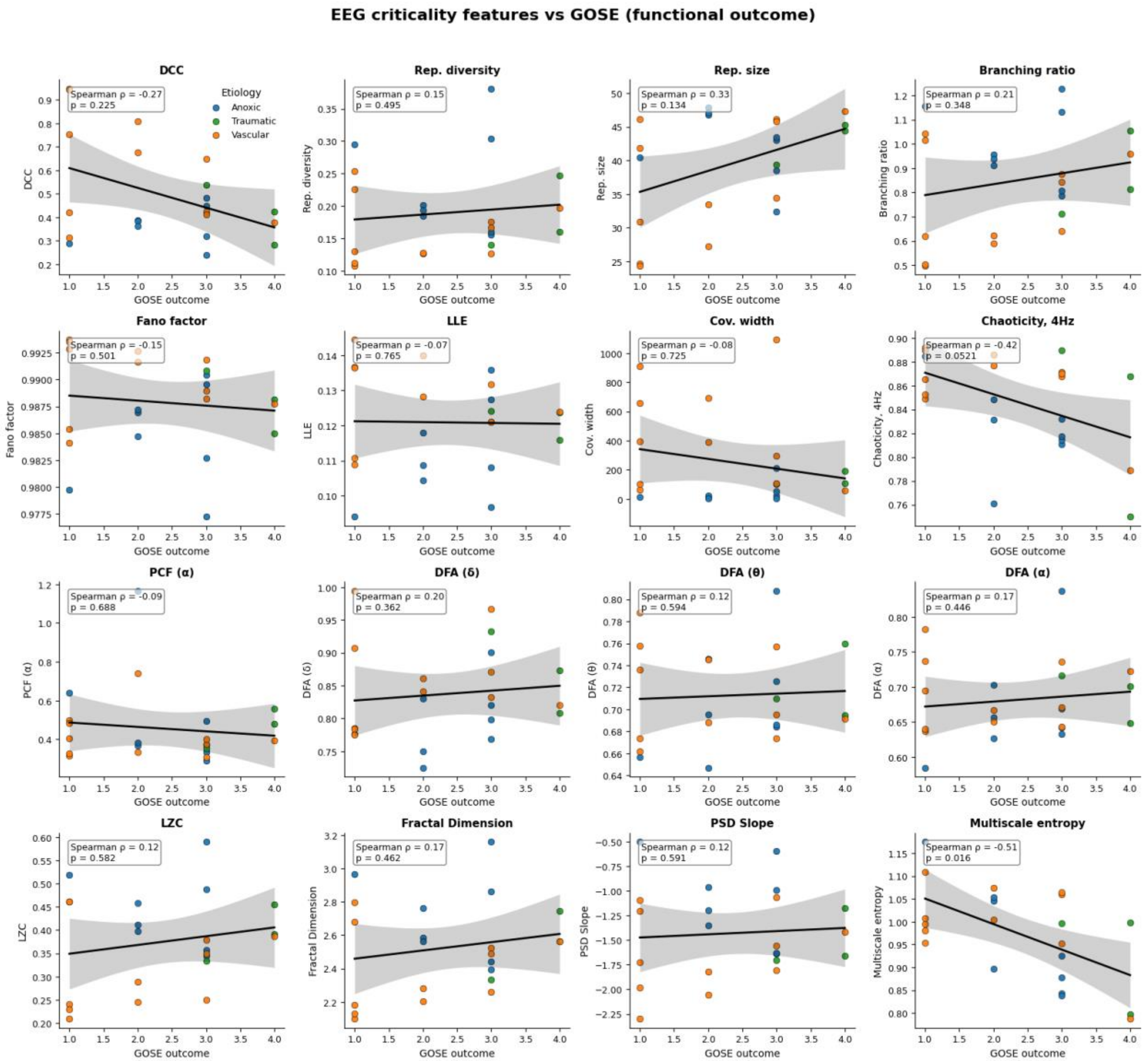


**Supplementary Figure 6. Associations between individual EEG criticality features and functional outcome (GOS-E).** Each panel shows the relationship between one resting-state EEG criticality feature and functional outcome as measured by the Glasgow Outcome Scale – Extended (GOS-E). Points represent individual patients and are colored by etiology (anoxic, vascular, traumatic). Black lines indicate fitted linear trend lines and gray shading denotes 95% confidence intervals. Each plot reports Spearman rank correlation coefficients and associated p-values.
